# Conserved GYXLI motif of FlhA is involved in dynamic domain motions of FlhA required for flagellar protein export

**DOI:** 10.1101/2022.03.25.485897

**Authors:** Tohru Minamino, Miki Kinoshita, Yumi Inoue, Akio Kitao, Keiichi Namba

## Abstract

Flagellar structural subunits are transported via the flagellar type III secretion system (fT3SS) and assemble at the distal end of the growing flagellar structure. The C-terminal cytoplasmic domain of FlhA (FlhA_C_) serves as a docking platform for export substrates and flagellar chaperones and plays an important role in hierarchical protein targeting and export. FlhA_C_ consists of domains D1, D2, D3, and D4 and adopts open and closed conformations. Gly-368 of *Salmonella* FlhA is located within the highly conserved GYXLI motif and is critical for the dynamic domain motions of FlhA_C_. However, it remains unclear how it works. Here, we report that periodic conformational changes of the GYXLI motif induce a remodeling of hydrophobic side-chain interaction networks in FlhA_C_ and promotes the cyclic open-close domain motions of FlhA_C_. The temperature- sensitive *flhA(G368C)* mutation stabilized a completely closed conformation at 42°C through strong hydrophobic interactions between Gln-498 of domain D1 and Pro-667 of domain D4 and between Phe-459 of domain D2 and Pro-646 of domain D4, thereby inhibiting flagellar protein export by the fT3SS. Its intragenic suppressor mutations reorganized the hydrophobic interaction networks in the closed FlhA_C_ structure, restoring the protein export activity of the fT3SS to a significant degree. Furthermore, the conformational flexibility of the GYXLI motif was critical for flagellar protein export. We propose that the conserved GYXLI motif acts as a structural switch to induce the dynamic domain motions of FlhA_C_ required for efficient and rapid protein export by the fT3SS.

**IMPORTANCE:** Many motile bacteria employ the flagellar type III secretion system (fT3SS) to construct flagella beyond the cytoplasmic membrane. The C-terminal cytoplasmic domain of FlhA (FlhA_C_), a transmembrane subunit of the fT3SS, provides binding-sites for export substrates and flagellar export chaperones to coordinate flagellar protein export with assembly. FlhA_C_ undergoes cyclic open-close domain motions. The highly conserved Gly-368 residue of FlhA is postulated to be critical for dynamic domain motions of FlhA_C_. However, it remains unknown how it works. Here, we carried out mutational analysis of FlhA_C_ combined with molecular dynamics simulation and provide evidence that the conformational flexibility of FlhA_C_ by Gly-368 is important for remodeling hydrophobic side-chain interaction networks in FlhA_C_ to facilitate its cyclic open-close domain motions, allowing the fT3SS to transport flagellar structural subunits for efficient and rapid flagellar assembly.

## INTRODUCTION

Many motile bacteria utilize flagella to swim in viscous liquids and move around on solid surfaces to migrate towards more favorable environments for their survival. The flagellum of *Salmonella enterica* serovar Typhimurium (hereafter referred to as *Salmonella*) is divided into three structural parts: the basal body, which acts as a bi- directional rotary motor; the filament, which functions as a helical propeller to produce thrust; and the hook, which is a universal joint connecting the basal body and filament and transmits torque produced by the motor to the filament. Flagellar assembly begins with the basal body, followed by the hook and finally the filament. To construct the flagellum on the cell surface, the flagellar type III secretion system (fT3SS) transports flagellar structural subunits from the cytoplasm to the distal end of the growing flagellar structure (1).

The fT3SS is located at the base of the flagellum and is composed of a transmembrane export gate complex with a stoichiometry of 9 FlhA, 1 FlhB, 5 FliP, 4 FliQ, and 1 FliR, and a cytoplasmic ATPase ring complex with a stoichiometry of 12 FliH, 6 FliI, and 1 FliJ (Fig. S1A in the Supplemental material). In addition, FlgN, FliS, and FliT act as export chaperones that escort their cognate substrates from the cytoplasm to the fT3SS (2, 3). The export gate complex utilizes the transmembrane electrochemical gradient of protons (H^+^) as the energy source to unfold and translocate export substrates across the cytoplasmic membrane (4, 5). The export gate complex requires ATP hydrolysis by the cytoplasmic ATPase ring complex to become an active H^+^/protein antiporter that couples H^+^ flow with protein translocation (6–8). The fT3SS also has a Na^+^-powered backup engine to maintain the flagellar assembly process when the cytoplasmic ATPase ring complex does not function properly (9, 10).

*Salmonella* FlhA is composed of an N-terminal transmembrane domain with eight transmembrane helices (FlhA_TM_, residues 1–327), a compactly folded cytoplasmic domain (FlhA_C_, residues 362–692) and a flexible linker (FlhA_L_, residues 328–361) connecting these two domains (11, 12) (Fig. S1B). FlhA_TM_ acts as a dual-ion channel that conducts both H^+^ and Na^+^. An interaction between FlhA_L_ and FliJ activates the ion channel of FlhA_TM_, allowing the export gate complex to couple either H^+^ or Na^+^ flow with the translocation of export substrate across the cytoplasmic membrane (13). FlhA_C_ forms a homo-nonamer in the fT3SS (14). The FlhA_C_ ring serves as a docking platform for export substrates and flagellar chaperones along with the C-terminal cytoplasmic domain of FlhB and plays an important role in hierarchical protein targeting and export for efficient flagellar assembly (15–21).

FlhA_C_ consists of domains D1, D2, D3, and D4 (Fig. S1B) and adopts open and closed conformations (12, 22). A highly conserved hydrophobic dimple is located at the interface between domains D1 and D2 and is directly involved in substrate recognition (16, 17). Because a large cleft exits in the interface between domains D2 and D4 in the open form of FlhA_C_, but not in the closed form (Fig. S1C), the chaperone-substrate complexes can bind to the conserved hydrophobic dimple in the open form but not in the closed form (23, 24). For filament assembly, the C-terminal region of FlhA_L_ binds to domains D1 and D2 of its neighboring FlhA_C_ subunit in the FlhA_C_ ring structure to stabilize the open conformation (FigS1C, right panel), allowing the chaperones- substrate complexes to efficiently bind to the FlhA_C_ ring (25). During hook assembly, however, FlhA_L_ binds to the conserved hydrophobic dimple of the open form not only to suppress premature docking of the chaperone-substrate complexes to FlhA_C_ but also to facilitate the export of the hook protein FlgE (26). These observations suggest that the open form of FlhA_C_ reflects an active state of the fT3SS. However, little is known about the role of the closed form of FlhA_C_ in flagellar protein export.

The *flhA(G368C)* mutation inhibits the protein transport activity of the fT3SS at a restrictive temperature of 42°C but not at a permissive temperature of 30°C (27–29). The temperature shift-up from 30°C to 42°C arrests the export of flagellar proteins by the fT3SS with the *flhA(G368C)* mutation. Molecular dynamics (MD) simulation has shown that the *flhA(G368C)* mutation restricts cyclic open-close domain motions of FlhA_C_ at 42°C and stabilizes a completely closed conformation of FlhA_C_ (24). Gly-368 is located within the highly conserved GYXLI motif (Fig. 1A). This leads to a plausible hypothesis that the conserved GYXLI motif may be critical for such domain motions of FlhA_C_ coupled with flagellar protein export.

**FIG 1.**
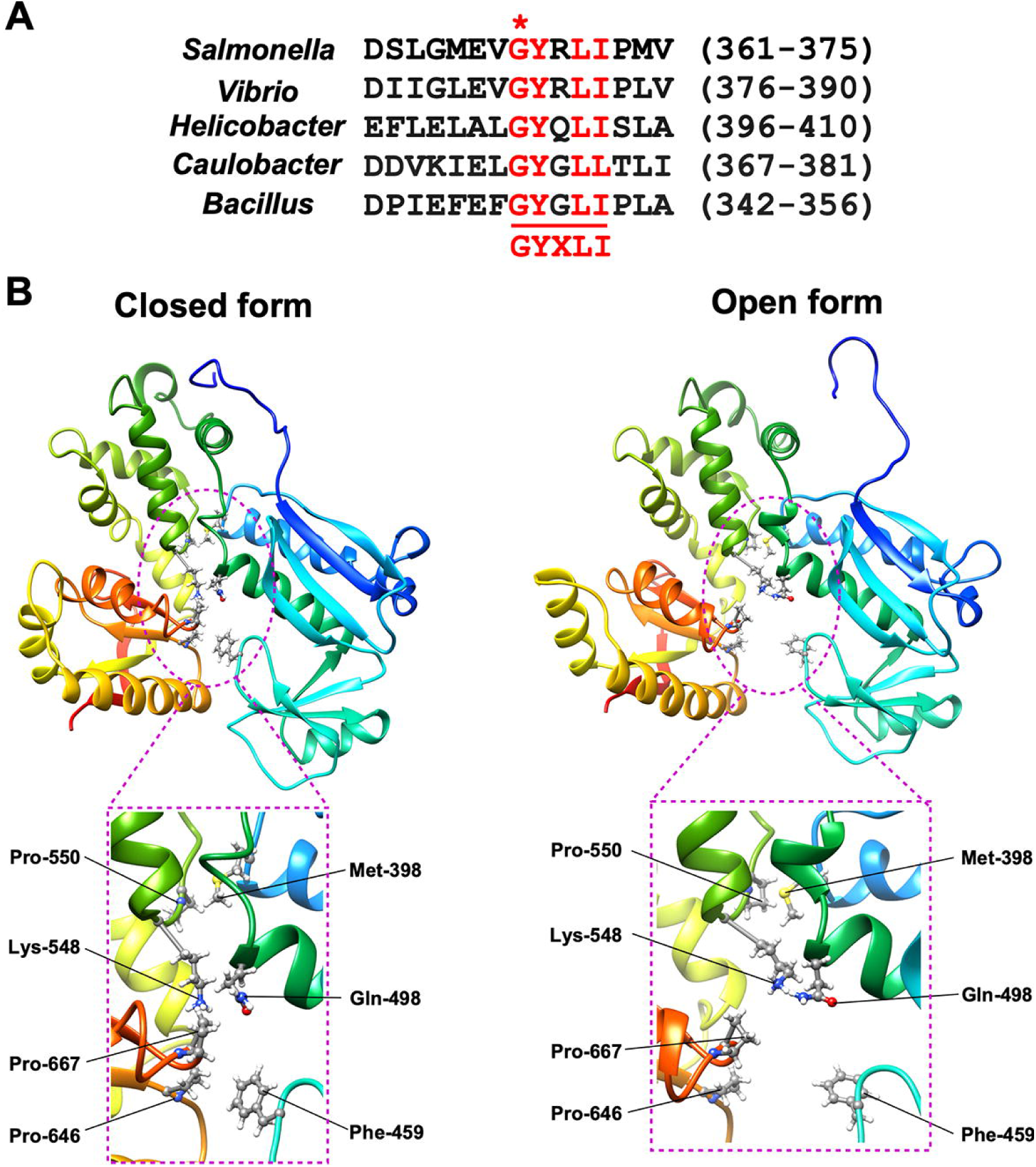
Role of a highly conserved Gly-368 residue of FlhA in dynamic domain motions of FlhA_C_. (A) Multiple sequence alignments of the conserved GYXLI motif of FlhA homologs. Multiple sequence alignment was carried out by Clustal Omega. Conserved residues are highlighted in red. A highly conserved glycine residue is indicated by an asterisk. UniProt Accession numbers: *Salmonella*, P40729; *Vibrio*, A0A6F8WT44; *Helicobacter*, O06758, *Caulobacter*, Q03845; *Bacillus*, P35620. (B) Cα ribbon diagrams of the closed (left panel) and open (right panel) forms of FlhA_C-G368C_ obtained by MD simulation. Phe-459 and Gln-498 make hydrophobic contacts with Pro-646 and Pro-667, respectively, in the closed from but not in the open form. The Cα backbone is color-coded from blue to red, going through the rainbow colors from the N- to the C-terminus.

To clarify this hypothesis, we carried out mutational analysis of FlhA_C_ combined with MD simulation. We provide evidence that the *flhA(G368C)* mutation stabilizes hydrophobic side-chain interactions between domains D1 and D3 and those between domains D2 and D4, thereby suppressing dynamic open-close domain motions of FlhA_C_, and that its intragenic suppressor mutations induce the remodeling of the hydrophobic interaction networks in FlhA_C_, allowing FlhA_C_ with the G368C mutation (FlhA_C-G368C_) to restore the dynamic open-close domain motions.

## RESULTS

### Effect of *flhA(G368C)* mutation on hydrophobic side-chain interaction networks in FlhA_C_

To address why the *flhA(G368C)* mutation stabilizes a completely closed form of FlhA_C_ at 42°C, we first compared the interfaces between domains D1 and D3 and those between domains D2 and D4 in the open and closed forms of FlhA_C-G368C_ obtained by MD simulation (Fig. 1B). Met-398 and Gln-498 of domain D1 make hydrophobic contacts with Pro-550 of domain D3 and Pro-667 of domain D4, respectively (left panel). Phe-459 of domain D2 makes a hydrophobic contact with Pro- 646 of domain D4 (left panel). Because the hydrophobic interactions between Phe-459 and Pro-646 and those between Gln-498 and Pro-667 are not seen in the open form of FlhA_C-G368C_ (right panel), we propose that the *flhA(G368C)* mutation may stabilize these two hydrophobic interactions at 42°C, thereby suppressing cyclic open-close motions of FlhA_C_.

### MD simulation of FlhA_C_ with either G368C/K548C or G368C/F459C/K548C mutation

The highly conserved Lys-548 residue of FlhA is located at the interface between domains D1 and D3 and contributes to D1-D3 interactions (12) (Fig. 1B). The *flhA(K548C)* mutation in domain D3 does not affect FlhA function at all. However, the combination of *flhA(G368C)* and *flhA(K548C)* mutations results in a loss-of-function phenotype even at 30°C (24). To clarify why the *flhA(G368C/K548C)* mutation inhibits flagellar protein export at 30°C, we analyzed the structure and dynamics of FlhA_C- G368C/K548C_ by MD simulation at 27°C for 1.5 μs. Domains D1 and D2 behaved as a rigid body in all cases. In contrast, domain D4 became closer to domain D2 through conformational changes of two hinges, one between domains D1 and D3 and the other between domains D3 and D4, when FlhA_C_ switched from an open conformation to a closed conformation. Therefore, we calculated the center-of-mass distance (*d*_24_) between domains D2 and D4 during MD simulation (Fig. 2A and Fig. S2 in the Supplemental material).

**FIG 2.**
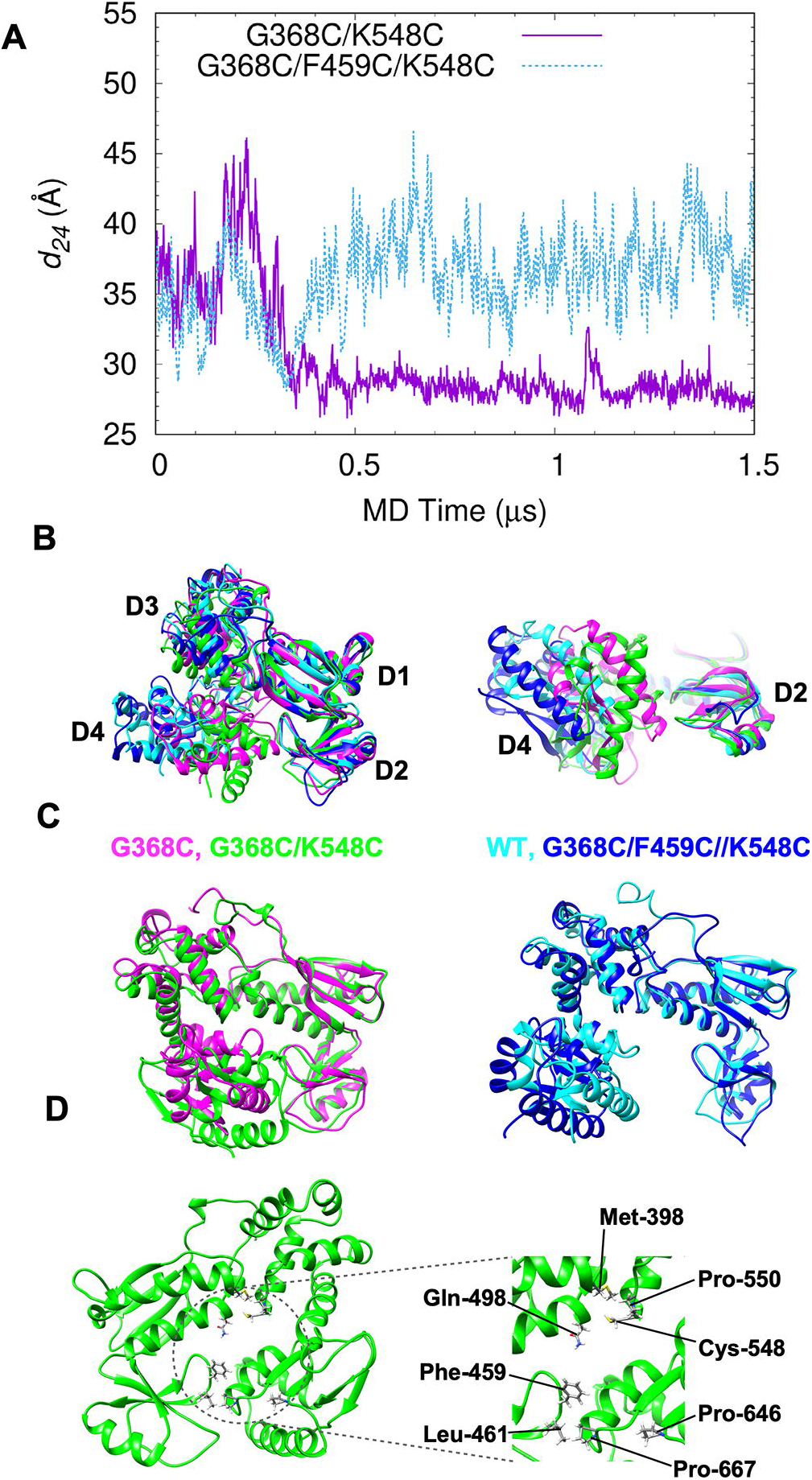
MD simulation of FlhA_C_ with either G368C/K548C or G368C/F459C/K548C. (A) Center-of-mass distance between domains D2 and D4 (*d_24_*) during 1.5 μs MD simulation. (B) Representative structures of wild-type (blue), FlhA_C-G368C_ (magenta), FlhA_C-G368C/K548C_ (green), and FlhA_C-G368C/F459C/K548C_ (cyan) obtained by MD simulation. Domains D1 and D2 are superimposed. (C) Structural comparisons between completely closed forms of FlhA_C-G368C_ (magenta) and FlhA_C-G368C/K548C_ (green) (left panels) and between the open forms of wild-type FlhA_C_ (blue) and FlhA_C-G368C/F459C/K548C_ (cyan). (D) Interfaces between domains D1 and D3 and between domains D2 and D4 of FlhA_C-G368C/K548C_

Unlike FlhA_C-G368C_, which changes its conformation back and forth between the closed and open forms at 27°C (24), FlhA_C-G368C/K548C_ stopped the dynamic domain motions at 27°C after 0.3 μs and switched its conformation to a completely closed form (Fig. 2A). When we compared the fully closed form of FlhA_C-G368C/K548C_ with that of FlhA_C-G368C_, there was a significant difference in the orientation of domain D4 relative to domain D3 (Fig. 2B and 2C, left panel). Unlike FlhA_C-G368C_, in which hydrophobic interactions between Gln-498 and Pro-667 and those between Phe-459 and Pro-646 stabilize the closed conformation of FlhA_C-G368C_ (Fig. 1B, left panel), the hydrophobic interactions of Pro-667 with Phe-459 and Leu-461 locked the FlhA_C-G368C/K548C_ structure into the fully closed conformation at 30°C (Fig. 2D).

It has been reported that the *flhA(F459C)* mutation restores motility of the *flhA(G368C/K548C)* mutant to a considerable degree (24), we analyzed the effect of this mutation on the structure and dynamics of FlhA_C-G368C/K548C_ by MD simulation. The F459C substitution restored the dynamic open-close domain motions of FlhA_C- G368C/K548C_ at 27°C (Fig. 2A). FlhA_C-G368C/F459C/K548C_ also adopted a super-open conformation (Fig. 2C, right panel) as seen in the wild-type structure at 42°C (24). Therefore, we conclude that the cyclic open-close domain motion of FlhA_C_ is required for efficient export of flagellar structural subunits by the fT3SS and that the completely closed form of FlhA_C_ reflects an inactive state of the fT3SS.

### Role of the highly conserved GYXLI motif for flagellar protein export by the fT3SS

We found structural differences in domain D1 when the open and closed forms of FlhA_C-G368C_ were compared (Fig. S3 in the Supplemental material). Gly-368 forms the highly conserved GYXLI motif along with Tyr-369, Arg-370, Leu-371, and Ile-372 (Fig. 1A), which forms a short α-helix (Fig. S3). Cys-368 of FlhA_C-G368C_ made much stronger hydrophobic contacts with Arg-370, Leu-413 and Pro-415 of domain D1 in the completely closed form than in its open form (Fig. 3A). As a result, the G368C mutation induced a significant conformational change of this α-helix (Fig. S3).

**FIG 3.**
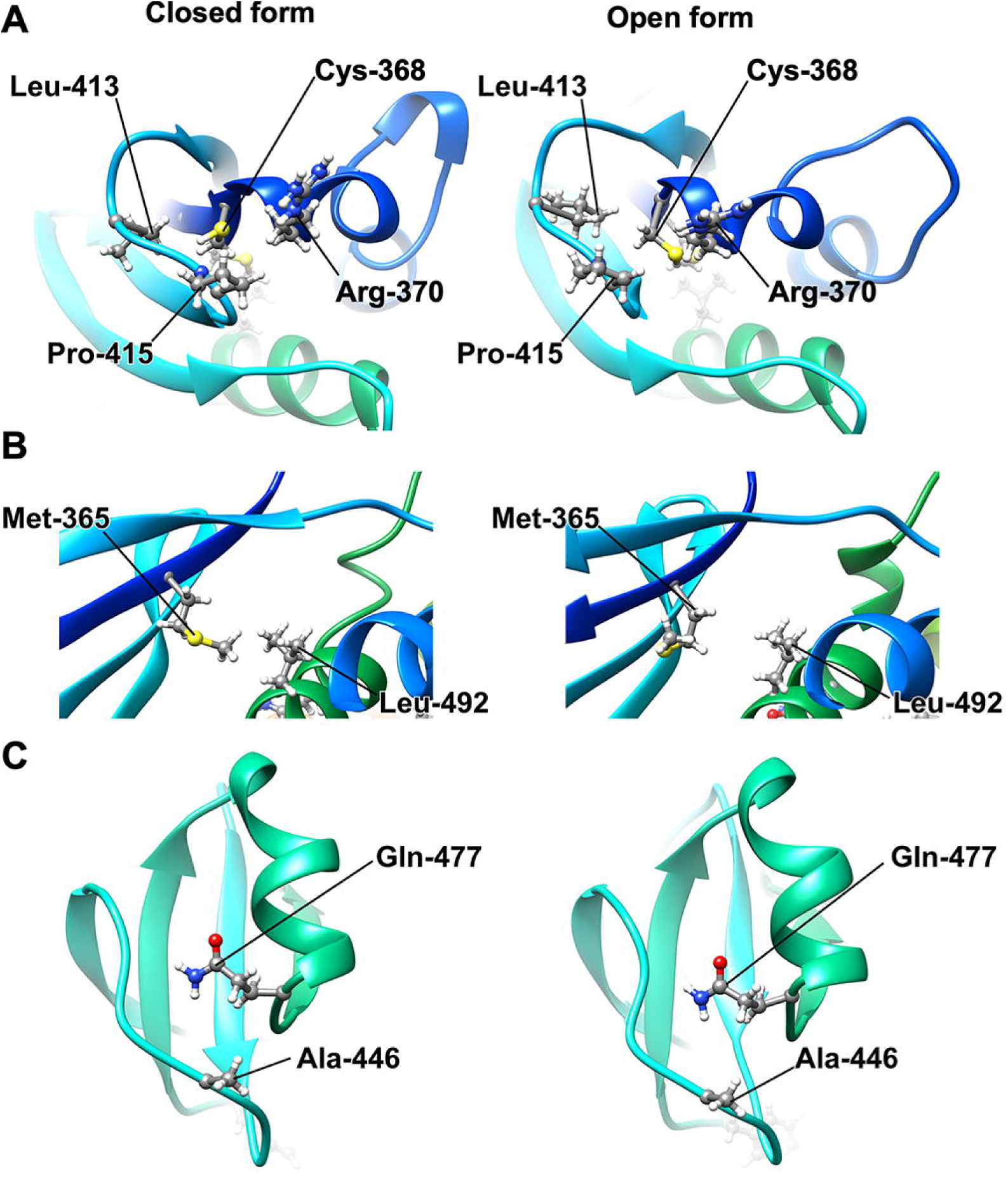
Structural comparison between closed (right panels) and open conformations (right panels) of FlhA_C-G368C_ obtained by MD simulation. (A) Hydrophobic interactions of Cys-368 with Arg-370, Leu-413, and Pro-415 in the open and closed forms of FlhA_C-G368C_. (B) Hydrophobic interactions between Met-365 and Leu-492. (C) Interaction between Gln-477 and Ala-446.

The *flhA(R370S)* mutation, which has been isolated as an intragenic suppressor mutation of the *flhA(G368C)* mutant grown at 42°C (28), is located within the conserved GYXLI motif (Fig. 1A). When the open and closed forms of FlhA_C-G368C_ obtained by MD simulation were compared, this R370S mutation presumably weakened the strong hydrophobic interactions among Cys-368, Leu-413, and Pro-415 (Fig. 3A), thereby partially restoring the protein export activity of the fT3SS with the G368C substitution. This raises a plausible hypothesis that that the conserved GYXLI motif may act as a structural switch to induce the cyclic open-close domain motion of FlhA_C_.

Hydrophobic parts of the side chain of Tyr-369 and Arg-370 make hydrophobic contacts with Pro-415 and Ala-416 in the open form of wild-type FlhA_C_, and Ile-372 stabilizes these hydrophobic interactions through the hydrophobic contact with Tyr-369. Leu-371 makes a hydrophobic contact with Pro-434 (Fig. 4A). To clarify the role of the conserved GYXLI motif of FlhA_C_ in flagellar protein export by the fT3SS, we constructed the Y369A/R370A/L371A/I372A (hereafter referred to as AAAA) and Y369G/R370G/L371G/I372G (hereafter referred to as GGGG) mutants and analyzed their motility in 0.35% soft agar. Immunoblotting with polyclonal anti-FlhA_C_ antibody revealed that the AAAA and GGGG mutations did not affect the steady expression level of FlhA (Fig. 4B). The AAAA mutant showed a very weak motility phenotype (Fig. 4B). In agreement with this motility phenotype, more than 80% cells of the AAAA strain produced shorter flagellar filaments compared to wild-type cells (Fig. 4C). The GGGG mutation caused a non-motile phenotype (Fig. 4B). Consistently, the GGGG mutant did not produce the flagellar filaments at all (Fig. 4C). These observations suggest that a conformational change of the GYXLI motif through remodeling of hydrophobic interaction networks seen in this motif may be critical for efficient flagellar protein export.

**FIG 4.**
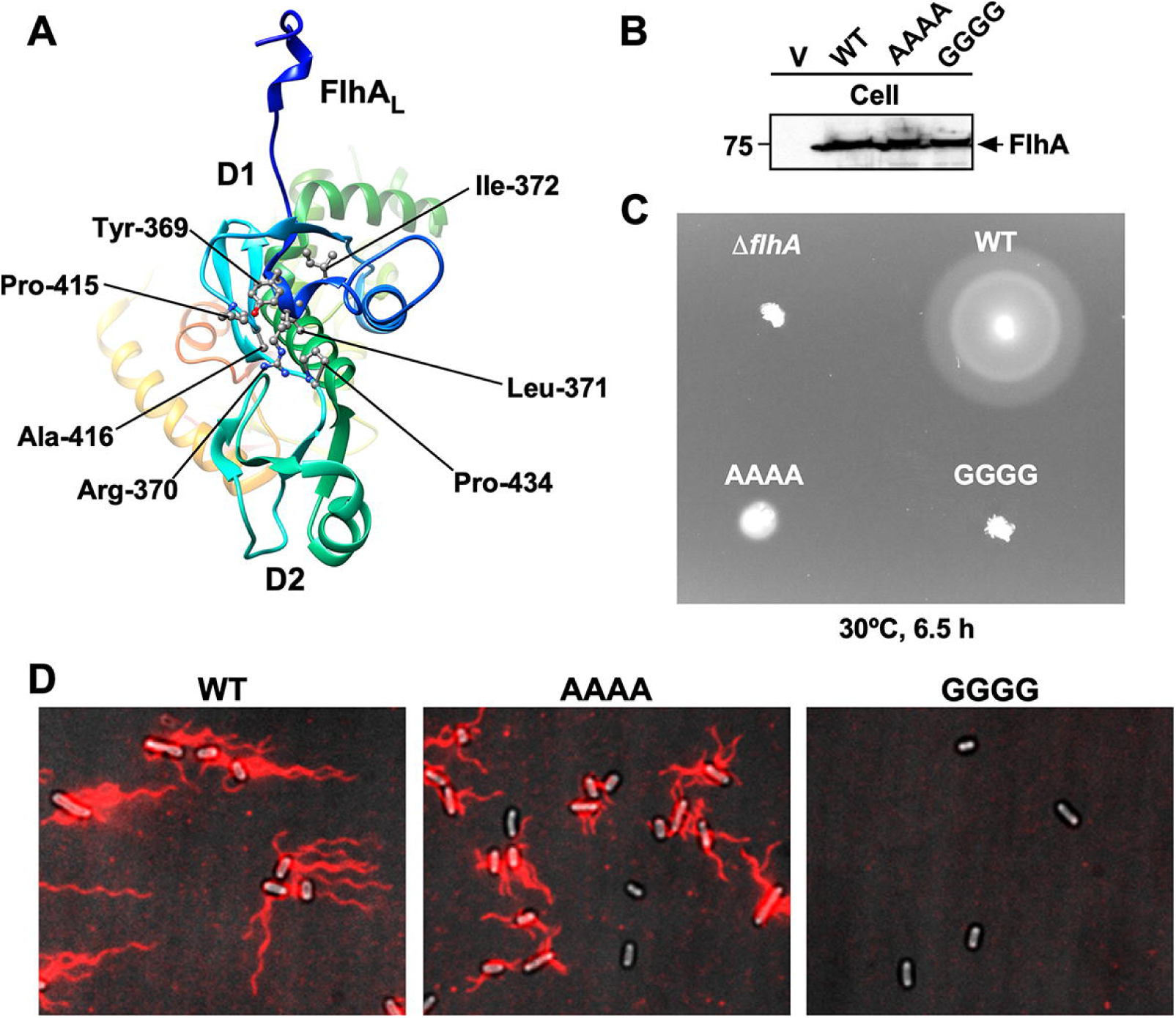
Effect of the Y369A/R370A/L371A/I372A and Y369G/R370G/L371G/I372G mutations on flagellar filament formation. (A) Hydrophobic side-chain network seen in the conserved GYXLI motif of wild-type FlhA_C_ (PDB ID: 3A5I). (B) Immunoblotting, using polyclonal anti- anti-FlhA_C_ antibody, of whole cell proteins (Cell) from NH001 (Δ*flhA*) carrying pTrc99AFF4 (indicated as V), pMM130 (indicated as WT), pMKM130- A4 (indicated as AAAA), and pMKM130-G4 (indicated as GGGG), which were exponentially grown at 30°C with shaking. (C) Motility of the above transformants in 0.35% soft agar. Plates were incubated at 30°C for 6.5 hours. (B) Fluorescent images of the same transformants. Fresh transformant cells were grown in L-broth containing ampicillin until the cells reached the stationary phase, and then flagellar filaments were labelled with a fluorescent dye, Alexa Fluor 594. The fluorescence images of the filaments labelled with Alexa Fluor 594 (magenta) were merged with the bright field images of the cell bodies.

### Effect of intragenic *flhA(M365I)*, *flhA(A446E)*, and *flhA(P550S)* suppressor mutations on the hydrophobic side-chain interaction networks

The intragenic *flhA(M365I)*, *flhA(A446E)*, and *flhA(P550S)* mutations have been reported to restore motility defects in the *flhA(G368C)* mutant at 42°C (24, 28). To clarify how the *flhA(G368C)* mutation stabilizes the completely closed form of FlhA_C_ at 42°C through the strong hydrophobic interactions between Gln-498 and Pro-667 and those between Phe-459 and Pro-646, we next analyzed the effect of the M365I, A446E, and P550S suppressor mutations on the conformational changes of FlhA_C-G368C_.

Pro-550 of domain D3 makes a hydrophobic contact with Met-398 of domain D1 (Fig. 1B), and therefore the P550S substitution should weaken the hydrophobic interactions between domains D1 and D3. Met-365 of domain D1 made a strong hydrophobic contact with Leu-492 of domain D1 in the closed form of FlhA_C-G368C_ but not in its open form (Fig. 3B). Therefore, we assume that the M365I substitution presumably affects this hydrophobic interaction to induce the conformational change of domain D1, thereby weakening the hydrophobic interaction between Gln-498 and Pro-667 in FlhA_C-G368C_.

It has been shown that the *flhA(G368C)* mutation induces a conformational change of domain D2 as judged by far-UV CD measurements of purified FlhA_C_ with the G368C substitution (29). Ala-446 of domain D2 hydrophobically interacted with Gln-477 of domain D2 in both open and closed forms of FlhA_C-G368C_ (Fig. 3C), and so the A446E mutation would induce the formation of a hydrogen bond between Glu-446 and Gln-447 as well as a hydrophobic contact. Therefore, we assume that this A446E mutation may induce a conformational change of domain D2, affecting the hydrophobic contact between Phe-459 and Pro-646.

### Mutational analysis of residues forming hydrophobic side-chain interaction networks in FlhA_C-G368C_

MD simulation of FlhA_C-G368C_ suggests that the *flhA(G368C)* mutation may induce hydrophobic side-chain interaction networks in FlhA_C_ to stabilize a complete closed form at 42°C and that Leu-413, Pro-415, Pro-434, Phe-459, Gln- 477, Leu-492, Gln-498, Lys-548, Pro-646, and Pro-667 may be involved in dynamic open-close domain motions of FlhA_C_ (Figs. 1B, 3, and 4A). To test whether the *flhA(G368C)* mutation affects the hydrophobic side-chain interaction networks in FlhA_C_, we constructed *flhA(G368C)* mutants with either L413A, P415A, P434A, F459A, Q477A, L492A, Q498A, K548A, P646A or P667A substitution and analyzed the motility of these double mutants at both 30°C and 42°C (Fig. 5A). These double mutations did not affect the steady expression level of FlhA as judged by immunoblotting with polyclonal anti-FlhA_C_ antibody (Fig. 5B).

**FIG 5.**
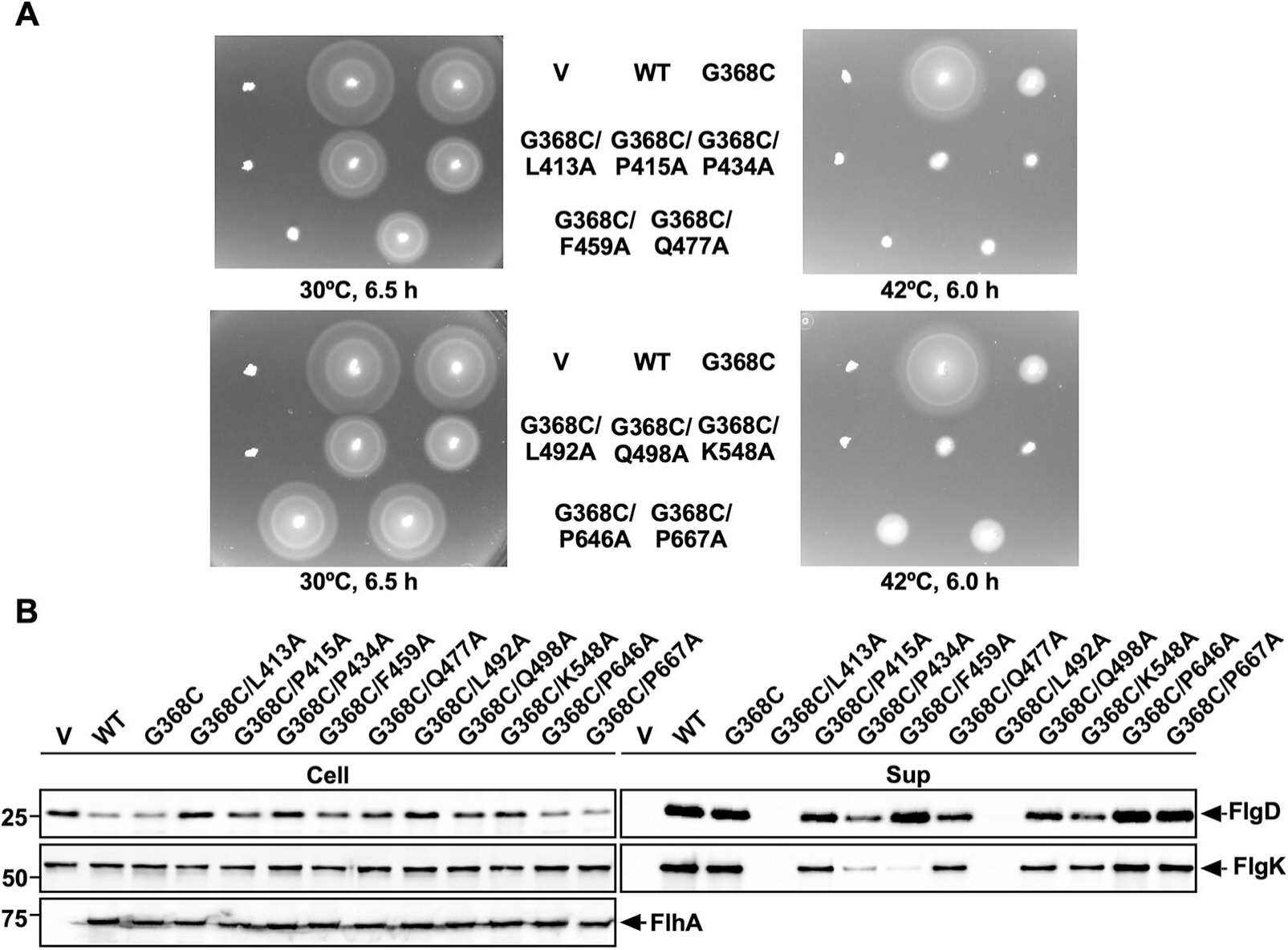
Effect of alanine substitution in residues involved in remodeling of hydrophobic side-chain interaction networks in FlhA_C-G368C_ on flagellar protein export by the *flhA(G368C)* mutant. (A) Motility of NH001 (Δ*flhA*) carrying pTrc99AFF4 (indicated as V), pMM130 (indicated as WT), pYI130(G368C) (indicated as G368C), pMKM130(G368C/L413A) (indicated as G368C/L413A), pMKM130(G368C/P415A) (indicated as G368C/P415A), pMKM130(G368C/P434A) (indicated as G368C/P434A), pMKM130(G368C/F459A) (indicated as G368C/F459A), pMKM130(G368C/Q477A) (indicated as G368C/Q477A), pMKM130(G368C/L492A) (indicated as G368C/L492A), pMKM130(G368C/Q498A) (indicated as G368C/Q498A), pMKM130(G368C/K548A) (indicated as G368C/K548A), pMKM130(G368C/P646A) (indicated as G368C/P646A) or pMKM130(G368C/P667A) (indicated as G368C/P667A) in 0.35% soft agar. Plates were incubated at 30°C for 6.5 hours and at 42°C for 6 hours. (B) Secretion assays of FlgD and FlgK. Immunoblotting, using polyclonal anti-FlgD (1st row), anti-FlgK (2nd row), or anti-FlhA_C_ (3rd row) antibody, of whole cell proteins (Cell) and culture supernatants (Sup) prepared from the above strains, which were exponentially grown at 30°C with shaking. The positions of FlgD, FlgK, and FlhA are indicated by arrows. Molecular mass markers (kDa) are shown on the left.

The P415A, P434A, Q477A, Q498A, and K548A substitutions reduced the motility of the *flhA(G368C)* mutant even at 30°C (Fig. 5A). Consistently, these five substitutions reduced the secretion levels of FlgD and FlgK by the fT3SS (Fig. 5B). Furthermore, they considerably reduced the motility of the *flhA(G368C)* mutant at 42°C (Fig. 5A). The P415A, P434A, Q477A, Q498A and K548A substitutions themselves displayed no phenotype at both 30°C and 42°C (Fig. 6A). Therefore, we suggest that the *flhA(G368C)* mutation affects the hydrophobic side-chain interaction networks in FlhA_C_, thereby inducing conformational changes of domains D1 and D2.

**FIG 6.**
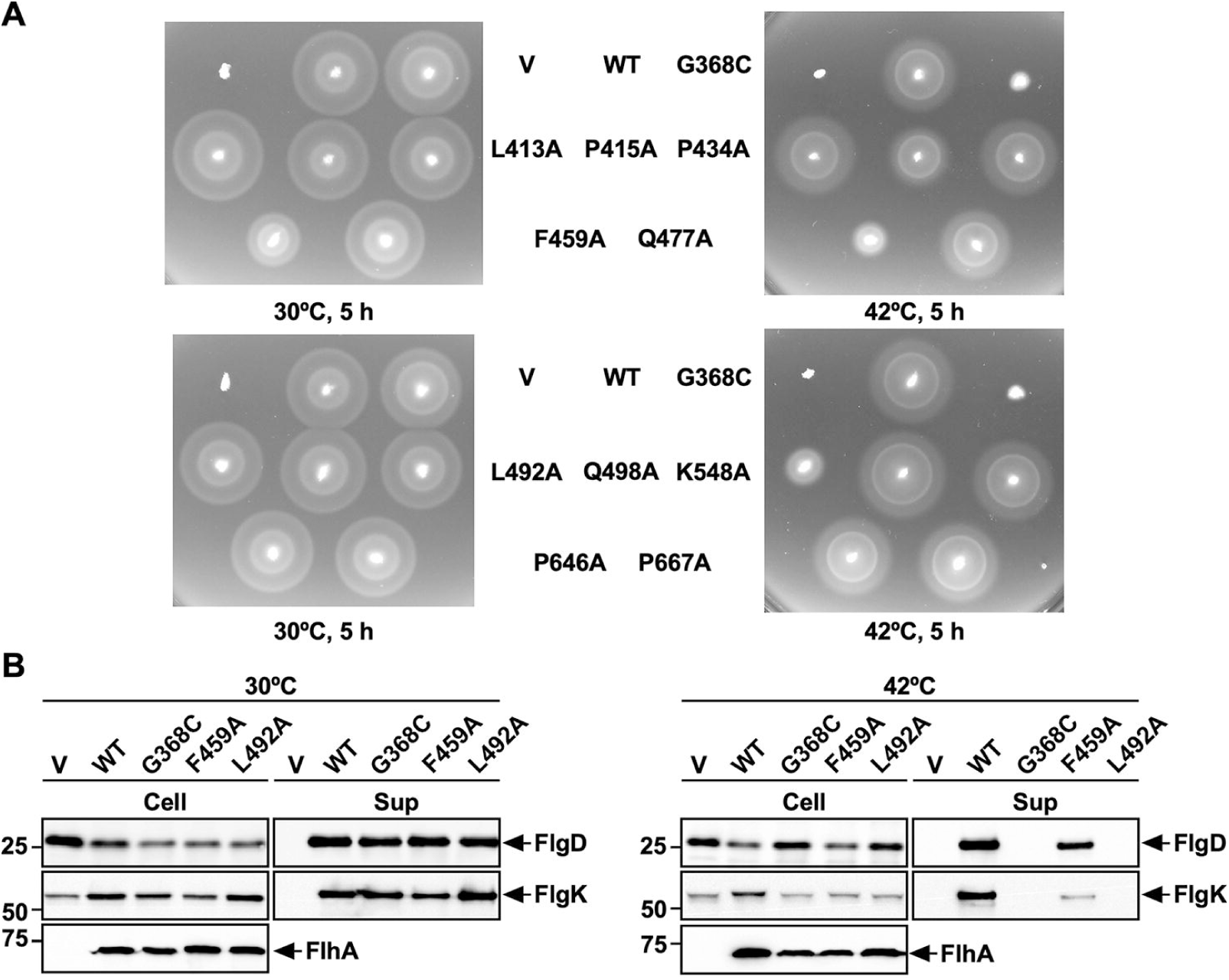
Effect of temperature on the protein export function of FlhA containing a single alanine substitution. (A) Motility of NH001 (Δ*flhA*) carrying pTrc99AFF4 (indicated as V), pMM130 (indicated as WT), pYI130(G368C) (indicated as G368C), pMKM130(L413A) (indicated as L413A), pMKM130(P415A) (indicated as P415A), pMKM130(P434A) (indicated as P434A), pMKM130(F459A) (indicated as F459A), pMKM130(Q477A) (indicated as Q477A), pMKM130(L492A) (indicated as L492A), pMKM130(Q498A) (indicated as Q498A), pMKM130(K548A) (indicated as K548A), pMKM130(P646A) (indicated as P646A)or pMKM130(P667A) (indicated as P667A) in 0.35% soft agar. Plates were incubated at 30°C and 42°C for 5 hours. (B) Secretion assays of FlgD and FlgK. Immunoblotting, using polyclonal anti-FlgD (1st row) or anti-FlgK (2nd row) antibody, of whole cell proteins (Cell) and culture supernatants (Sup) prepared from NH001 carrying pTrc99A, pMM130, pYI130(G368C), pMKM130(F459A), or pMKM130(L492A) which were exponentially grown at either 30°C (left panels) or 42°C (right panels). The positions of FlgD, and FlgK are indicated by arrows. Molecular mass markers (kDa) are shown on the left.

The L413A and L492A substitutions interfered with the motility of the *flhA(G368C)* mutant at both 30°C and 42°C (Fig. 5A). Consistently, these two substitutions inhibited the secretion of both FlgD and FlgK by the *flhA(G368C)* mutant grown at 30°C (Fig. 5B). The L413A mutation itself did not inhibit motility at either 30°C or 42°C (Fig. 6A, upper panels). Because Cys-368 makes strong hydrophobic contacts with Arg-370, Leu-413 and Pro-415 of domain D1 in the completely closed form of FlhA_C-G368C_ (Fig. 3A), we suggest that the L413A mutation stabilizes these hydrophobic contacts even at 30°C, thereby inhibiting the protein transport activity of the fT3SS. In contrast to the L413A mutation, the L492A substitution itself significantly reduced motility at 42°C but not at 30°C (Fig. 6A, lower panels). Consistently, this substitution inhibited the secretion of FlgD and FlgK by the fT3SS at 42°C but not at 30°C (Fig. 6B). These results indicate that the temperature shift-up from 30°C to 42°C induces a conformational change of domain D1 of FlhA_C_ by the L492A substitution, thereby reducing the protein export activity of the fT3SS. Because Leu-492 makes a hydrophobic contact with Met-365 in the closed form of FlhA_C-G368C_ but not in its open form (Fig. 3B), we suggest that a proper switching of the hydrophobic side-chain interaction between Met-365 and Leu-492 is critical for flagellar protein export by the fT3SS.

Phe-459 of FlhA is located within a highly conserved hydrophobic dimple at the interface between domains D1 and D2 (Fig. 1B) and is directly involved in substrate recognition (16,17,23,24). The F459A mutation reduced the motility of the *flhA(G368C)* mutant at both 30°C and 42°C (Fig. 5A, upper panels). This mutation reduced the level of FlgK secretion by the *flhA(G368C)* mutant grown at 30°C but not the level of FlgD secretion (Fig. 5B), indicating that the F459A substitution affects the docking of the FlgN-FlgK chaperone-export substrate complex to FlhA_C-G368C_ but not that of FlgD, in agreement with previous reports (17, 21). Because the motility of the *flhA(G368C/F459A)* was much worse than that of the *flhA(F459A)* mutant (Figs. 5 and 6), we suggest that the G368C mutation induces a conformational change of the conserved hydrophobic dimple involved in the interaction with flagellar chaperones. Furthermore, the temperature shift-up from 30°C to 42°C reduced the levels of both FlgD and FlgK secretion by the *flhA(F459A)* mutant (Fig. 6B), thereby reducing motility at 42°C (Fig. 6A). Therefore, we propose that a remodeling of hydrophobic side-chain interaction networks in the conserved hydrophobic dimple may be required for efficient and stable binding of export substrates and chaperone-substrate complexes to FlhA_C_. The P646A and P667A mutations, which are located within domain D4, did not affect the motility of the *flhA(G368C)* mutant at both 30°C and 42°C (Fig 3A, lower panels). These two mutations themselves showed no phenotype (Fig4A, lower panels), indicating that these two proline residues are not critical for the FlhA function. The structural transition from the open conformation to the closed conformation occurs in at least two steps (24). The first conformational change occurs at the interface between domains D3 and D4, resulting in a 13° rotation of domain D4 in the direction towards domain D2, and the second conformational change occurs at a flexible hinge between domains D1 and D3, allowing FlhA_C_ to adopt the closed form. Therefore, we suggest that the *flhA(G368C)* mutation may limit the hinge movement between domains D1 and D3 through a remodeling of the hydrophobic side-chain interaction networks of domain D1, thereby allowing Pro-646 and Pro-667 to make the strong hydrophobic contacts with Phe-459 and Gln-498, respectively (Fig. 1B).

## Discussion

The temperature-sensitive *flhA(G368C)* mutation limits the cyclic open-close domain motions of FlhA_C_ at 42°C, thereby reducing the flagellar protein export by the fT3SS and therefore the cell motility (24). Gly-368 is located within the highly conserved GYXLI motif of FlhA_C_ (Fig. 1A), suggesting that the GYXLI motif may be involved in such domain motions of FlhA_C_. To clarify the role of Gly-368 in flagellar protein export, we performed mutational analysis of FlhA_C-G368C_ combined with MD simulation and showed that the *flhA(G368C)* mutation induced a remodeling of hydrophobic side-chain interaction networks in FlhA_C_ at 42°C, allowing Gln-498 of domain D1 and Phe-459 of domain D2 to hydrophobically interact with Pro-667 and Pro-646 of domain D4, respectively (Fig. 1B). As a result, the *flhA(G368C)* mutation not only suppresses the dynamic open-close domain motions of FlhA_C_ but also stabilizes the completely closed conformation at 42°C. Intragenic M365I, R370S, A446E and P550S suppressor mutations, which restore motility defects in the *flhA(G368C)* mutant to near wild-type levels (24, 28), affected the hydrophobic side-chain interaction networks in the closed FlhA_C_ structure, thereby restoring the protein export activity of the fT3SS containing the *flhA(G368C)* mutation (Fig. 3). Cys-368 hydrophobically interacts with Arg-370, Leu-413 and Pro-415 of domain D1 in the closed form of FlhA_C-G368C_ but not in its open form (Fig. 3A). Because the AAAA and GGGG mutations reduced motility considerably (Fig. 4B), we propose that the conformational flexibility of FlhA_C_ by Gly-368 may be required for cyclic conformational changes of the conserved GYXLI motif, allowing FlhA_C_ to undergo the cyclic open-close domain motions through the remodeling of the hydrophobic side-chain interaction networks in FlhA_C_.

FlhA_C_ forms a nonameric ring structure in the fT3SS (14, 30). The cryo-EM structure of the SctV_C_ ring derived from the enteropathogenic *Escherichia coli* T3SS injectisome, which is a FlhA_C_ homolog, has shown that the SctV_C_ subunit adopts a closed conformation and maintained the closed conformation during MD simulation (31). In contrast, the cryo-EM structure of SctV_C_ of *Salmonella* SPI-2 T3SS injectisome undergoes dynamic open-closed domain motions in a way similar to FlhA_C_ (32). Here, we showed that the completely closed form of FlhA_C_ reflected an inactive state of the fT3SS and that the cyclic open-close domain motions of FlhA_C_ was critical for efficient and rapid flagellar protein transport by the fT3SS (Fig. 2). The FlgN, FliS, and FliT chaperones in complex with their cognate substrates bind to FlhA_C_ at a nanomolar affinity (17, 33), and such strong interactions between the chaperones and FlhA_C_ assist protein unfolding and export by the fT3SS (34, 35). The flagellar chaperones bind to the open form of FlhA_C_ but not to the closed form (23, 24). Purified FlhA_C-G368C_ prefers to adopt a closed conformation even at room temperature (24). Because pull-down assays by GST affinity chromatography have revealed that the *flhA(G368C)* mutation reduces the binding affinity of FlhA_C_ for the chaperone-substrate complex (24), we propose that the structural transition of FlhA_C_ from the open to the closed form may induce the dissociation of empty chaperone from FlhA_C_ for the binding of a newly delivered chaperone-substrate complex for the export of next substrate.

FliJ binds to FlhA_L_ with high affinity and opens both the H^+^ and polypeptide channels of the transmembrane export gate complex, allowing the fT3SS to couple H^+^ flow through the H^+^ channel with the translocation of export substrate through the polypeptide channel (Fig. S1B in the Supplemental material) (6,15,26). FliJ also binds to FlhA_C_ with low affinity (26). It has been shown that CdsO, a FliJ homolog, binds to CdsV_C_, a FlhA_C_ homolog, at a large cleft between the D4 domains of neighboring CdsV_C_ subunits in the CdsV_C_ ring structure (36). This suggests that FliJ also binds to the cleft between the D4 domains of neighboring FlhA_C_ subunits in the FlhA_C_ ring. It has been reported that FliJ stabilizes the interaction between FlhA_C_ and FliT in complex with FliD (15). This raises the possibility that the interaction of FliJ with the cleft between the D4 domains in the FlhA_C_ ring may stabilize the open conformation of FlhA_C_ subunits in the ring. To clarify this possibility, we built the models of the FlhA_C_ ring in complex with FliJ in the open and closed forms of FlhA_C_ based on the crystal structure of the CdsV_C_ ring in complex with CdsO (PDB ID: 6WA9) (Fig. 7). FliJ can bind to the cleft between the D4 domains of neighboring FlhA_C_ subunits stably when FlhA_C_ adopts the open form in the ring structure (right panel) because FliJ can contact both FlhA_C_ subunits. However, the FliJ binding to of FlhA_C_ in the closed form appears to be unstable (left panel) because the distance between the neighboring D4 domains is longer in the closed form than that in the open form, restraining the proper contacts of FliJ with two FlhA_C_ subunits (middle panel). Because the open and closed forms of FlhA_C_ represent the active and inactive states of the fT3SS, respectively (Fig. 2), we propose that the cyclic open-close domain motion of FlhA_C_ may be important for efficient and robust energy coupling mechanism of the fT3SS.

**FIG. 7.**
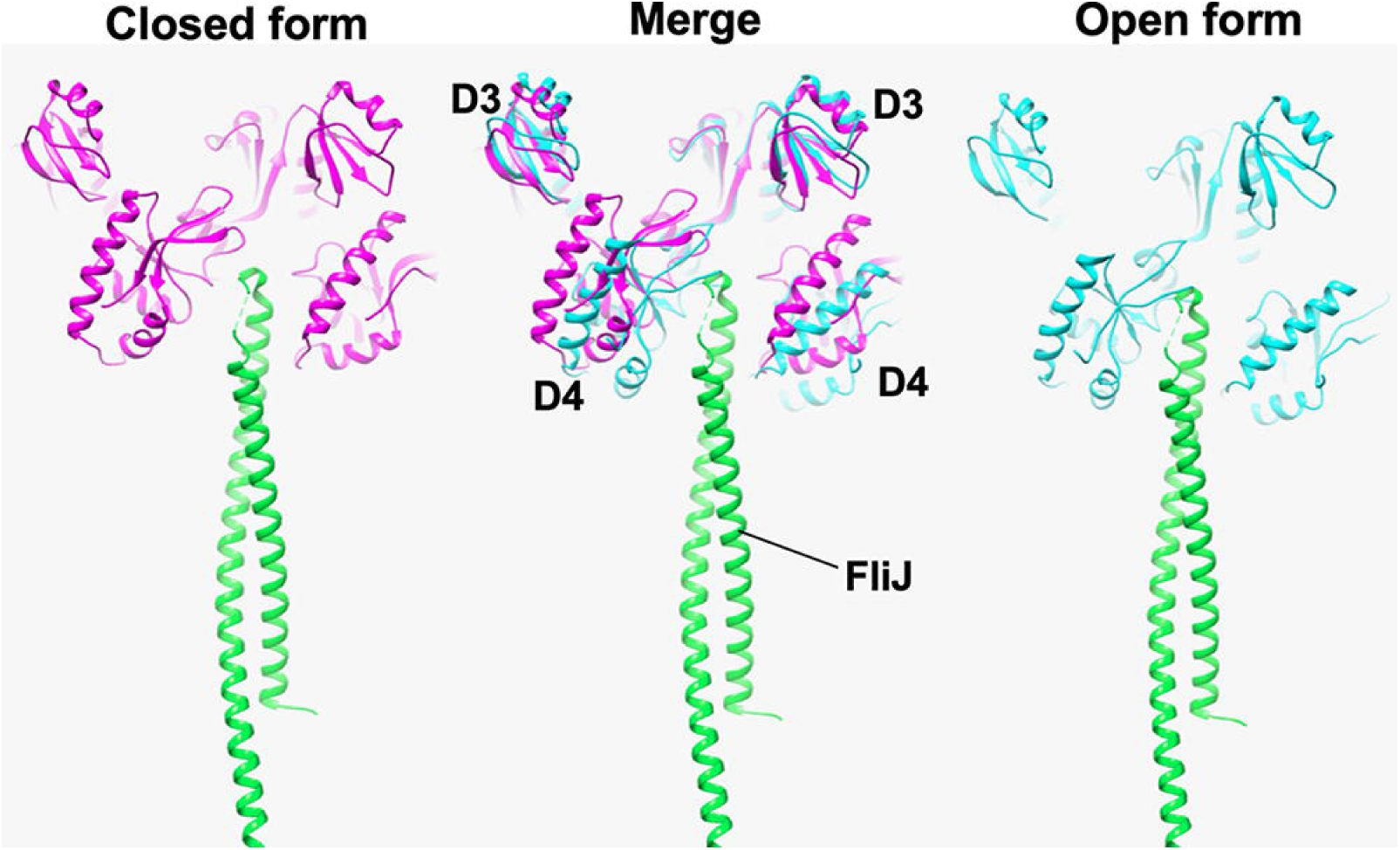
Model for of the closed (left panel, magenta) and open (right panel, cyan) forms of the FlhA_C_ ring with FliJ. The closed and open ring models were made by fitting domains D1 and D2 of the open form of FlhA_C_ (PDB ID: 3A5I) and its closed form obtained by MD simulation to those of MxiA_C_ in the nonameric ring structure (PDB ID: 4A5P). The CdsO-CdsV_C_ complex structure (PDB ID: 6WA9) was superimposed on the FlhA_C_ ring models, and then FliJ (green) (PDB ID: 3AJE) was superimposed on the 6WA9 structure to build the FlhA_C_-FliJ ring complex. Two FlhA_C_ subunits in the ring model are shown. FliJ can bind to a cleft between domains D4 of neighboring FlhA_C_ subunits in the open form of the FlhA_C_ ring (right panel). Because a distance between the D4 domains is longer in the closed form than in the open form (center panel), FliJ presumably cannot bind to the cleft in the closed form.

## MATERIALS AND METHODS

### Bacterial strains, P22-mediated transduction, and DNA manipulations

*Salmonella* strains used in this study are listed in Table 1. P22-mediated transductional crosses were performed with P22HT*int* (37). DNA manipulations were performed using standard protocols. Site-directed mutagenesis was carried out using Prime STAR Max Premix as described in the manufacturer’s instructions (Takara Bio). All of the *flhA* mutations were confirmed by DNA sequencing (Eurofins Genomics).

### Motility assay

Fresh colonies were inoculated onto soft agar plates [1% (w/v) triptone, 0.5% (w/v) NaCl, 0.35% (w/v) Bacto agar] containing 50 μg/ml ampicillin, and then the plates were incubated at 30°C. At least six measurements were performed.

### Observation of flagellar filaments with a fluorescent dye

*Salmonella* cells were grown at 30°C in 5 ml of L-broth [1% (w/v) Bacto-tryptone, 0.5% (w/v) Bacto-yeast extract, 0.5% (w/v) NaCl] containing 100 μg/ml ampicillin. The cells were attached to a cover slip (Matsunami glass, Japan), and unattached cells were washed away with motility buffer (10 mM potassium phosphate pH 7.0, 0.1 mM EDTA, 10 mM L-sodium lactate). A 1 μl aliquot of polyclonal anti-FliC serum was mixed with 50 μl of motility buffer and then 50 μl of the mixture was applied to the cells attached to the cover slip. After washing with the motility buffer, 1 μl of anti-rabbit IgG conjugated with Alexa Fluor 594 (Invitrogen) was added to 50 μl of motility medium, and then the mixture was applied. After washing with the motility buffer, the cells were observed by fluorescence microscopy (38). Fluorescence images were analyzed using ImageJ software version 1.52 (National Institutes of Health).

### Secretion assay

*Salmonella* cells were grown in 5 ml of L-broth containing 100 μg/ml ampicillin at either 30°C or 42°C with shaking until the cell density reached an OD_600_ of ca. 1.2–1.4. Cultures were centrifuged to obtain cell pellets and culture supernatants. The cell pellets were resuspended in sodium dodecyl sulfate (SDS)-loading buffer solution [62.5 mM Tris-HCl, pH 6.8, 2% (w/v) SDS, 10% (w/v) glycerol, 0.001% (w/v) bromophenol blue] containing 1 μl of 2-mercaptoethanol. Proteins in the culture supernatants were precipitated by 10% trichloroacetic acid and suspended in a Tris/SDS loading buffer (one volume of 1 M Tris, nine volumes of 1 X SDS-loading buffer solution) containing 1 μl of 2-mercaptoethanol (39). Both whole cellular proteins and culture supernatants were normalized to a cell density of each culture to give a constant number of *Salmonella* cells. After boiling proteins in both whole cellular and culture supernatant fractions at 95°C for 3 min, these protein samples were separated by SDS–polyacrylamide gel electrophoresis and transferred to nitrocellulose membranes (Cytiva) using a transblotting apparatus (Hoefer). Then, immunoblotting with polyclonal anti-FlgD, anti-FlgK, or anti-FlhA_C_ antibody was carried using iBind Flex Western Device (Thermo Fisher Scientific) as described in the manufacturer’s instructions. Detection was performed with Amersham ECL Prime western blotting detection reagent (Cytiva). Chemiluminescence signals were captured by a Luminoimage analyzer LAS-3000 (Cytiva). At least three measurements were performed.

### Multiple sequence alignment

Multiple sequence alignment was carried out using CLUSTAL-Ω (http://www.ebi.ac.uk/Tools/msa/clustalo/).

### MD simulation

MD simulations of FlhA_C_ with either G368C/K548C or G368C/F459C/K548C substitution were conducted as described previously (24). The mutant structures were constructed based on the 3A5I structure of FlhA_C_ (12). AMBER ff14SB force field (40) was used for the proteins. Each mutant was initially solvated in a cubic box of SPC/E_b_ water molecules (41) and 0.17 M KCl (42) with a margin of at least 12 Å from the proteins to the periodic box boundaries. The simulation was conducted with the *pmemd.cuda* module (43) of AMBER20/21 (44). MD simulations of the G368C/K548C and G368C/F459C/K548C mutants were independently conducted at 300 K. After energy minimization and equilibration MD for 1 ns with positional restraints for the protein main chain atoms at 300 K and 1 atm, MD simulation without restraints were conducted for 1.5 μs.

## Supporting information

Sipplrmrntal Figures

## ACKNOWLEDGEMENTS

This work was supported in part by JSPS KAKENHI Grant Numbers JP20K15749 and 22K06162 (to M.K.), JP19H03191, JP20H05439, and JP21H05510 (to A.K.), and JP19H03182 and 22H02573 (to T.M.). This work has also been supported by Platform Project for Supporting Drug Discovery and Life Science Research (BINDS) from AMED under Grant Number JP19am0101117 to K.N., by the Cyclic Innovation for Clinical Empowerment (CiCLE) from AMED under Grant Number JP17pc0101020 to K.N. and by JEOL YOKOGUSHI Research Alliance Laboratories of Osaka University to K.N.

## Notes

### Competing Interest Statement

The authors have declared no competing interest.

### Summary of Updates

We revised our manuscript based on reviewers' comments. We added one additional data set to the revised manuscript and also modified all figures.

