## Supplementary material for "Conserved GYXLI motif of FlhA is involved in dynamic domain motions of FlhA required for flagellar protein export": Sipplrmrntal Figures

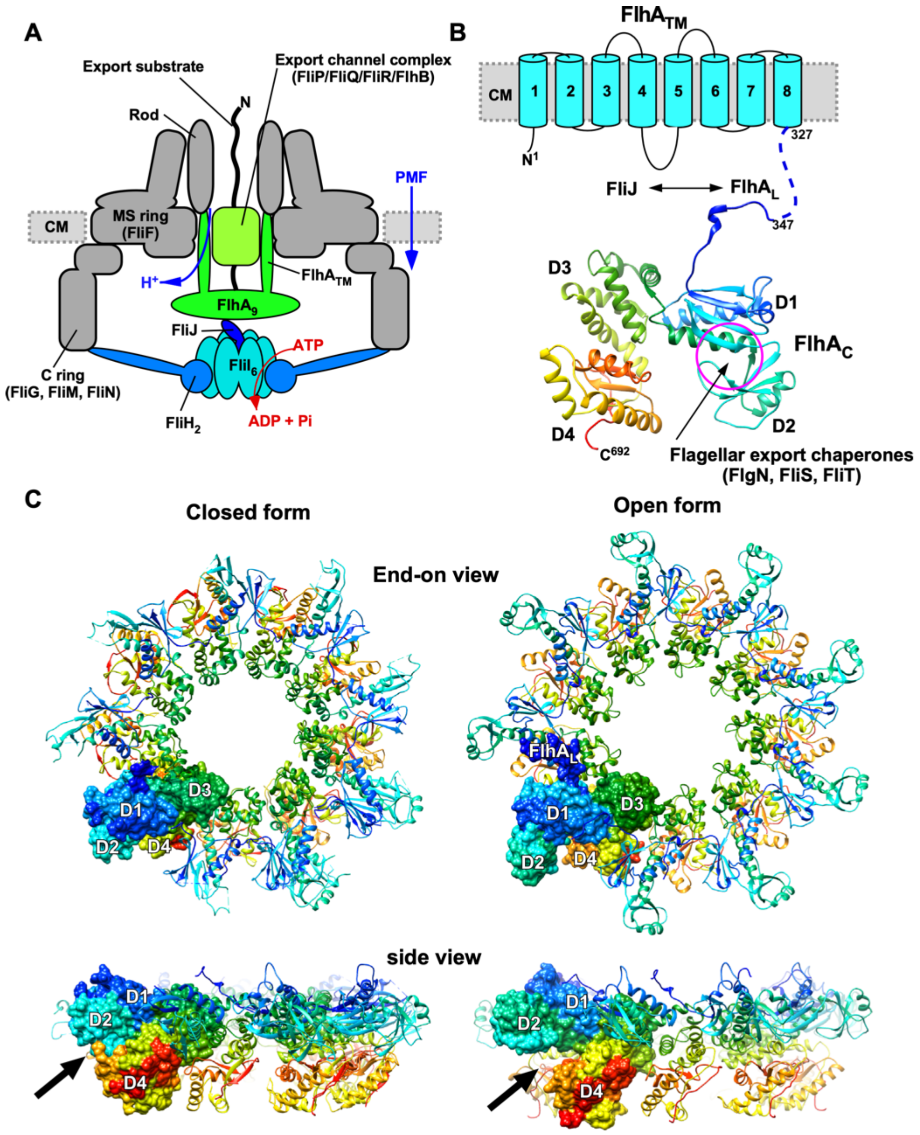


**FIG S1. Flagellar type III secretion system (fT3SS).** (A) Schematic diagram of the fT3SS. The fT3SS is composed of a transmembrane export gate complex made of five transmembrane proteins, FlhA, FlhB, FliP, FliQ, and FliR, and a cytoplasmic ATPase ring complex consisting of three soluble proteins, FliH, FliI, and FliJ. The export gate complex is located inside the MS ring and utilizes proton motive force (PMF) across the cytoplasmic membrane (CM) to drive proton ($H^+$)-coupled protein export. FlhA forms a homo-nonamer through interactions between the C-terminal cytoplasmic domain of FlhA ($FlhA_C$), and its N-terminal transmembrane domain ($FlhA_{TM}$) acts as a transmembrane ion channel that conduct both $H^+$ and $Na^+$. The cytoplasmic ATPase ring complex associates with the C ring through an interaction between FliH and FliN. ATP hydrolysis by the ATPase ring complex activates the export gate complex through an interaction between $FlhA_L$ and FliJ, allowing the gate complex to become an active protein transporter that couples $H^+$ flow through the FlhA proton channel to protein translocation. (B) Topological model of FlhA. FlhA is composed of an N-terminal transmembrane domain ($FlhA_{TM}$) with eight transmembrane helices (1–8), a large C-terminal cytoplasmic domain ($FlhA_C$), and a flexible linker connecting $FlhA_{TM}$ and $FlhA_C$.

$FlhA_C$ (PDB ID: 3A5I) has four domains (D1, D2, D3, and D4). The Cα backbone is color-coded from blue to red, going through the rainbow colors from the N- to the C-terminus. Flagellar export chaperones in complex with their cognate substrates bind to a conserved hydrophobic dimple at the interface between domains D1 and D2. (C) Open and closed $FlhA_C$ ring models. The$FlhA_C$ ring models were made by fitting domains D1 and D2 of the open form of $FlhA_C$ (PDB ID: 3A5I) and its closed form obtained by MD simulation to those of $MxiA_C$ in the nonameric ring structure (PDB ID: 4A5P). Arrows indicate flagellar chaperone binding sites in the ring model.

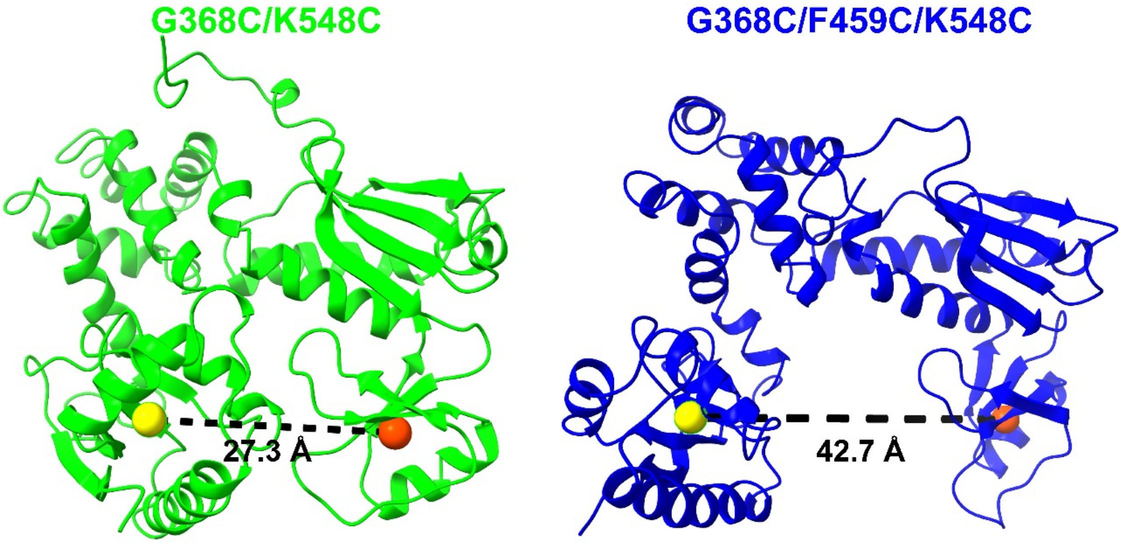


**FIG S2. Examples of center-of-mass distance between domains D2 and D4 (*$d_{24}$*) for $FlhA_{C\text{-}G368C/K548C}$ and $FlhA_{C\text{-}G368C/F459C/K548C}$.** The $d_{24}$ distances of $FlhA_{C\text{-}G368C/K548C}$ (indicated as G368C/K548C) and $FlhA_{C\text{-}G368C/F459C/K548C}$ (indicated as G368C/F459C/K548C) at 1.5 μs and 300 K are shown. Orange and yellow spheres indicate the center-of-mass positions of D2 and D4, respectively. Each image was created by Chimera X (doi: 10.1002/pro.3235).

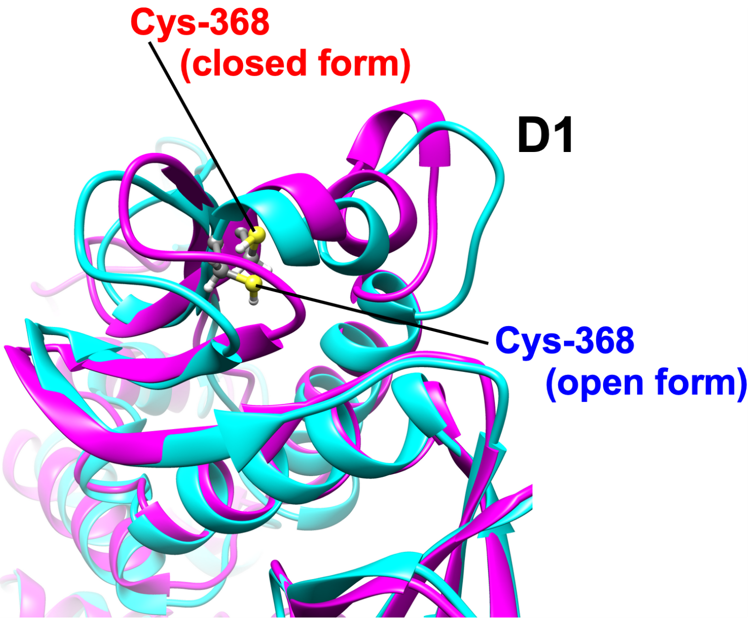


**FIG S3. Structural comparison between the closed (magenta) and open (cyan) forms of FlhA$_{C\text{-}G368C}$ obtained by MD simulation.** The temperature shift-up from 30ºC to 42ºC induces a conformational change of domain D1 through remodeling of hydrophobic side-chain interaction networks in FlhA$_{C\text{-}G368C}$.
